# Minipuberty regulates reproductive lifespan and ovarian follicular loss in a mouse model with reduced minipubertal gonadotropin levels

**DOI:** 10.1101/2024.08.20.608775

**Authors:** Mélanie Chester, Marie M Devillers, Raphaël Corre, Frank Giton, Fatoumata Souaré, Claire-Hélène Petrovic, Éloïse Airaud, Daniel Quintas, Sakina Mhaouty-Kodja, Lydie Naulé, Céline J Guigon

**Author notes:** Correspondence address: Dr Céline Julie Guigon, Université Paris Cité, Inserm, 3PHM, Pathophysiology and Pharmacotoxicology of the Human Placenta, pre & postnatal Microbiota, Paris, F-75006, France.

## Abstract

**Study question:** What is the role of the physiological hypergonadotropic activity encountered at minipuberty on the implementation of female reproductive function, in a mouse model with manipulated minipubertal gonadotropin levels?

**Summary answer:** Elevated minipubertal levels of gonadotropins may have long-term effects on fertility by mediating neuroendocrine aging and ovarian follicle depletion.

**What is known already:** Minipuberty is characterized by the tremendous activation of the gonadotropin axis, as evidenced by elevated levels of gonadotropins regulating folliculogenesis as well as the synthesis of ovarian hormones including estradiol, testosterone, and AMH.

**Study design, size, duration:** To determine whether hypergonadotropic activity of the gonadotropin axis at mini-puberty could impact reproductive parameters and female fertility, we used a pharmacological approach to suppress gonadotropin levels in Swiss mice by injecting daily a GnRH receptor antagonist (GnRHR) (Ganirelix, 10 μg/mouse) or its vehicle between 10 and 16 postnatal days, to cover the entire duration of minipuberty. We analyzed the onset of puberty and estrous cyclicity as well as fertility in young (3 to 5 months) and middle-aged (11 months) mice from control (CTR) and antagonist-treated groups (n = 17 to 20 mice/age and treatment group). Ovaries and brains were collected, fixed and sectioned (for histology, follicle count and immunohistochemistry) or frozen (for analysis of follicular markers, aging and inflammation) from adult females, and blood was collected by cardiac puncture for hormonal assays (n = 3 to 8 mice/age and treatment group).

**Participants/materials, setting, methods:** To analyze the initiation of puberty, we monitored vaginal opening and performed vaginal smears to detect first estrus and diestrus 2 in control and antagonist-treated mice. We studied estrous cyclicity on vaginal smears to detect the occurrence of the different stages of the cycle at the beginning of reproductive life. Young and middle-aged mice of the two groups were mated several times with males to assess fertility rates, delay of conception and litter size. To evaluate ovarian function, we counted follicles at the primordial, primary, secondary and tertiary stages and corpora lutea by morphometric analyses, and we determined the relative intra-ovarian abundance of follicular markers (*Amh*, *Inhba*, *Inhbb*, *Cyp19a1*, *Lhcgr*, *Fshr*) by real-time RT-PCR, as well as the levels of circulating AMH and progesterone by ELISA and GC/MS, respectively. We also analyzed features of ovarian aging and inflammation (presence of oocyte-depleted follicles and multinucleated giant cells) by histology and by measuring the relative intra-ovarian abundance of *Sirt1*, *Sod2*, *Tnfa* and *Il1b* using real-time RT-PCR. To determine the impact on neuroendocrine determinants related to the control of reproduction, we analyzed circulating gonadotropin levels using Luminex assays as well as kisspeptin and GnRH immunoreactivity by immunohistochemistry in the hypothalamus, in both young and middle-aged mice.

**Main results and the role of chance:** Our results show that the treatment had no impact on the initiation of puberty, estrous cyclicity, or fertility at the beginning of reproductive life. However, it increased reproductive lifespan, as shown by the higher percentage of antagonist-treated females than controls (33% versus 6%) still fertile at 11 months of age (*P*=0.0471). There were no significant differences in the number of kisspeptin and GnRH neurons, nor in the density of kisspeptin- and GnRH-immunoreactivity in the hypothalamic areas involved in reproduction between the two groups of mice studied at either 4 or 11 months. In addition, basal levels of LH and FSH were comparable between the two groups at 4 months, but not those of LH at 11 months which were much lower in females treated with antagonist than in their age-matched controls (237 ± 59.60 pg/mL in antagonist-treated females versus 1027 ± 226.3 pg/mL in controls, *P*=0.0069). Importantly, at this age, antagonist-treated mice had basal LH levels comparable to young mice (e.g., in 4-month-old controls: 294 ± 71.75 pg/mL, P > 0.05), while those of control females were higher (*P*= 0.0091). Despite their prolonged reproductive lifespan and delayed neuroendocrine aging, antagonist-treated mice exhibited earlier depletion of their follicles, as shown by lower numbers of primordial, primary, and secondary follicles associated with lower circulating AMH levels and relative intra-ovarian abundance of *Amh* transcripts than control mice. However, they exhibited comparable completion of folliculogenesis, as suggested by the numbers of tertiary follicles and corpora lutea, relative intra-ovarian abundance of *Cyp19a1*, *Inha* and *Inhb* transcripts, and circulating progesterone levels that all remained similar to those of the control group. These observed alterations in ovarian function were not associated with increased ovarian aging or inflammation.

**Large scale data:** none

**Limitations, reasons for caution:** This study was carried out on mice, which is a validated research model. However, human research is needed for further validation.

**Wider implications of the findings:** This study, which is the first to investigate the physiological role of minipuberty on reproductive parameters, supports the idea that high postnatal levels of gonadotropins may have long-term effects on female fertility by regulating the duration of reproductive life. Changes in gonadotropin levels during this period of life, such as those observed in infants born prematurely, may thus have profound consequences on late reproductive functions.

**Study funding/competing interest(s):** This research was conducted with the financial support of ANR AAPG2020 (ReproFUN), CNRS, Inserm, Université Paris Cité and Sorbonne Université. The authors declare that they have no conflicts of interest.

## Introduction

In females, fertility involves a tight dialogue between the different components of the gonadotrope axis during the reproductive cycle, i.e., the hypothalamus, the pituitary and the ovary. The pulsatile release of GnRH drives the circulating levels of gonadotropins, acting on the ovary to regulate follicular growth and ovulation, as well as the production of hormones such as estradiol (E2), which in turn adjusts the activity of the hypothalamus and pituitary (Richards and Pangas, 2010). Before the advent of the cyclic activity of the gonadotrope axis at puberty, there is a short and tremendous activation of the gonadotrope axis during minipuberty of infancy. This phase, occurring shortly after birth (10-17 days postnatal in mice; 1 week-6 months in girls) (Winter *et al*., 1975; François *et al*., 2017), corresponds to high levels of gonadotropin hormones and significant production of different hormones including E2, testosterone and AMH (Penny *et al*., 1974; Winter *et al*., 1975; Burger *et al*., 1991; François *et al*., 2017; Devillers *et al*., 2019, 2023). Observations in humans and other mammal species support the idea of a possible action of minipubertally-produced E2 on the growth of mammary gland and uterus (Devillers *et al*., 2022). However, one can assume that this period could also play a role in the maturation of the gonadotrope axis and, thus, in fertility. Indeed, hypothalamic networks involved in GnRH pulsatility, including kisspeptin neurons of the rostral periventricular nucleus of the third ventricle area (RP3V) which are involved in triggering puberty and ovulation in response to the positive feedback exerted by E2, are still maturing during this period (Clarkson and Herbison, 2006; Clarkson *et al*., 2009). In addition, the ovary is already host to gonadotropin-dependent regulation of sex steroids and AMH by the first waves of primary and preantral/early antral follicles, subsequently contributing to ovulations up to the very beginning of reproductive life (Guigon *et al*., 2003; Zheng *et al*., 2014; François *et al*., 2017; Devillers *et al*., 2019, 2023). There is also a high rate of recruitment of primordial follicles which will contribute to ovulations much later in adult life (Zheng *et al*., 2014). We hypothesized that the particular endocrine environment of minipuberty could have short- and long-term effects on reproductive function by acting on the maturation of the gonadotrope axis.

To determine whether minipuberty regulates key reproductive parameters, i.e., the onset of puberty, estrous cyclicity and fertility, we adopted a pharmacological approach to suppress gonadotropin levels in mice by injecting a GnRH receptor (GnRHR) antagonist during the entire duration of minipuberty. The current study clearly demonstrates that the treatment had no significant effects on puberty initiation, estrous cyclicity and early reproductive life. However, this resulted in prolonged fertility of middle-aged mice associated with basal LH levels similar to those of reproductively active young mice, despite accelerated follicular depletion. Our study highlights a possible regulatory role of minipuberty in reproductive senescence.

## Materials and methods

### Animals and treatments

Studies were conducted on SWISS mice aged 10-16 days post-natal (dpn), adult cycling (1 to 5 months) and middle-aged females (11 months) that were born at the animal facility from genitors purchased at Janvier Labs (Le Genest St Isle, France). The day the pups were born was considered as day 0 dpn of age and animals were weaned at 21 dpn. Mouse care and handling was performed as previously described (François *et al*., 2017). Ten µg of GnRH antagonist (Ganirelix, Fyremadel Gé, Ferring S.A.S, France) or saline were subcutaneously injected daily between 10 and 16 dpn. Mice were anesthetized with a mix of ketamine (Imalgene® 1000) and xylazine (Rompun® 2%) to collect blood by cardiac puncture. Blood was allowed to clot at room temperature for at least 15 minutes, and then centrifuged at 5000 g for 5 minutes to obtain the serum. After cervical dislocation, ovaries were collected, frozen in liquid nitrogen and stored at −80 °C for RNA extraction, or fixed in Bouin’s solution (HT10132, Sigma Aldrich) for histological analysis. Experiments were performed in accordance with standard ethics guidelines and were approved by the Institutional Animal care and Use committee of the University Paris Cité by the French Ministry of Agriculture (CEB#13-2021).

### Determination of circulating gonadotropin levels

LH and FSH were assayed in 10 μl of serum using the Luminex technology with the rat pituitary magnetic bead panel Milliplex Map kit (Merck-Millipore, Nottingham, UK) in accordance with the manufacturer’s instructions, as described previously (François *et al*., 2017). The sensitivity of the assays was 32 pg/mL for FSH and 3.2 pg/mL for LH. The inter-assay coefficient of variation was 5.4% for FSH and 3.2% for LH. The intra-assay coefficient of variation was 7.6% for FSH and 5.9% for LH.

### Determination of estrous cyclicity

Estrous cyclicity was analyzed by vaginal smears performed daily for approximately 10 days following vaginal opening in peripubertal mice and for 21 days in 14–18-week-old females. Smears were taken from mice between 9:30 and 10:30 a.m. using a saline solution (NaCl 0.9%) and analyzed after staining with hematoxylin/eosin (HE). The stages of the estrous cycle were characterized by the predominance of nucleated cells, keratinized (or cornified) cells and leukocytes for proestrus (PE), estrus (E), diestrus 1 and diestrus 2 (D2), respectively.

### Fertility tests

Twenty control (CTR) and 19 antagonist-treated females aged 3 months-old were mated with wild-type adult males (2 females with 1 male/cage). Body weight was monitored at the beginning of the mating and every week. Females that gained at least 4 g of weight in the week following mating were isolated until delivery, which was monitored daily. Once the number of pups was recorded at birth (litter size), pups were killed and females were placed with different males for additional mating (at approximatively 4 and 5 months-old, and at 11 months-old). Time to conception was considered as the time between the day of mating and the day of delivery.

### Immunofluorescence and microscopy analysis

Females at 4 and 11 months in diestrus were anesthetized with a mix of ketamine (Imalgene® 1000) and xylazine (Rompun® 2%) before being perfused with phosphate buffer (PB) (0.1M; pH=7.4) and paraformaldehyde (PFA) 4%. Brains were collected and post-fixed for 24 hours in 4% PFA-PB solution then cryoprotected in 30% sucrose-PB solution. Coronal sections (30 µm) were sliced using a cryostat Leica CM3050S and processed for kisspeptin/GnRH dual-label immunofluorescence as previously described (Naulé *et al*., 2015, 2023). Sections were blocked for 2h with 2% normal donkey serum (Sigma-Aldrich) in phosphate buffer saline (PBS)-Triton 0.3%, then incubated with sheep anti-kisspeptin (1:5000, INRAE #AC053) and rabbit anti-GnRH #19900 (dilution: 1:2000;(Caldani *et al*., 1988), Histochemistry) for 72h at 4°C. Immunofluorescence was performed using Alexa Fluor 488-conjugated donkey anti-sheep (1:500, Invitrogen #A11015) and Alexa Fluor 555-conjugated donkey anti-rabbit (1:500, Invitrogen #A21429) secondary antibodies. Brain sections were dried and mounted with Mowiol. Using a Zeiss A320 microscope, the number of GnRH-immunoreactive neurons were examined on two anatomically matched sections selected at the level of the rostral preoptic area (rPOA) (plates 26-27 of the Mouse Brain Atlas of Paxinos and Franklin). The number of kisspeptin cells was counted in the anteroventral periventricular (AVPV), rostral (rPeN) and caudal (cPeN) preoptic periventricular nuclei (plates 28 to 30, two sections per animal). Using a Zeiss LSM 710 confocal microscope, GnRH fiber density was quantified by voxel counts at the level of the median eminence (ME) (plate 45, two sections per animal) and the total kisspeptin-immunoreactivity (ir) was carried out in the medial region of the arcuate nucleus (ARC) (plate 45, two sections per animal) as previously described (Naulé *et al*., 2015, 2023).

### Tissue processing for histological analyses

For histological analyses, at least 6 to 7 ovaries of 11 months-old mice treated with either Ganirelix or its vehicle were fixed for 48 hours in Bouin’s solution, rinsed in phosphate buffer solution (PBS) and placed in ethanol 70%. The ovaries were then dehydrated and paraffin-embedded (HISTIM core facility, Cochin Institute, Paris, France). Sections of 5 µm thickness were mounted on glass slides at the frequency of one section in five, and they were stained with hematoxylin and eosin, using routine procedures. After dehydration in alcohol and mounting in Eukitt (Sigma), slides were scanned using Pannoramic 250 Flash III, 3D HISTECH (Cellular and tissue imaging core facility, Bicêtre Biomedical Institute). Classification of primordial, primary, secondary (preantral), tertiary (antral) follicles and oocyte-depleted follicles (ODF) was previously described (Guigon *et al*., 2003, 2005). Follicular counting was performed on every five tissue sections. To facilitate the analysis, a Deep Learning segmentation program was developed based on MMdetection tools https://github.com/open-mmlab/mmdetection using a resnet 50 FastRCNN pretrained model implemented with a mask detection layer. The model was initially trained on 3 manually segmented ovaries and segmentation results were uploaded on a databank Omero server https://github.com/ome/omero-server to manually correct and complement the detections. To avoid repeated counting of the same preantral/antral follicles on several tissue sections, when a follicle was seen for the first time, it was marked and tracked in each subsequent section throughout which it appeared. Only follicles with visible oocytes were counted.

### Determination of serum progesterone levels

The mass spectrometry coupled with gas chromatography (GC-MS) procedure was used to measure progesterone concentrations in 11-months old mice ovaries, as described previously (Giton *et al*., 2015; Marie *et al*., 2023). The linearity of steroid measurement was confirmed by plotting the ratio of the steroid peak response/internal standard peak response to the concentration of steroid for each calibration standard. Accuracy, precursor ion analyte, corresponding deuterated internal control, range of detection, low limit of quantification (LLOQ), and intra & inter assay CVs of the quality control are reported in Supplementary table 1.

### Determination of serum AMH levels

Quantification of AMH in the serum was performed using the commercial rat and mouse AMH ELISA kit (Ansh Labs, AL-113) following the manufacturer’s protocol, as previously described (Devillers *et al*., 2019). Samples were diluted at 1/10 and light absorbance was read at 450 nm using a microplate reader (FlexStation, Molecular device). Limit of ELISA sensitivity was 0.34 ng/mL. The intra- and inter-assay variability was 1.6+/- 0.53% and 5.7+/-1.6% respectively.

### RNA extraction, reverse transcription, and quantitative real-time PCR

Single frozen ovaries were lysed with TissueLyser II (Qiagen) and RNA was obtained using RNeasy mini-kit (#74106 Qiagen), following manufacturer’s protocol, as previously described (François *et al*., 2017). Total RNA (1 µg) was reverse transcribed using random primers and Superscript II (Invitrogen) following manufacturer’s instructions. Primers used for quantitative real-time PCR are listed in Table 1. Real-time PCR was performed with LightCycler 480 SYBR Green I Master and LightCycler 480 instrument (Roche Molecular Biochemicals, La Rochelle, France), as previously described (Devillers *et al*., 2019). Experiments were performed using ovarian samples from CTR and antagonist-treated mice, with each sample run in triplicates.

**Table 1.**
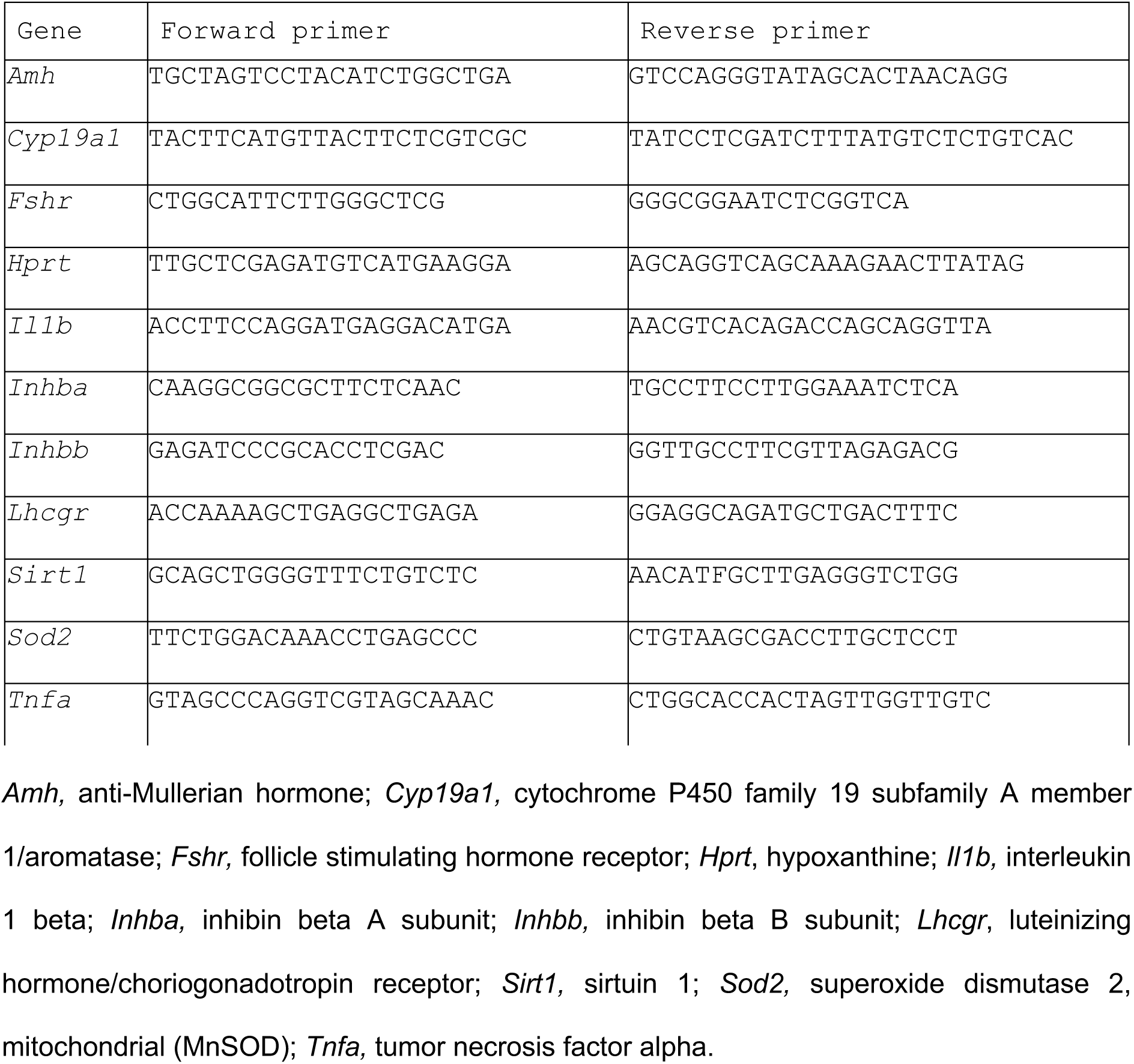
Primers used for quantitative RT-PCR analysis.

### Statistical analysis

For each experiment, the number of samples per age/treatment group is indicated in the legend of the figure. Data were analyzed using Prism 9 (version 9.0, GraphPad Software). Depending on experimental setting, statistical analyses were performed by Student t-test (one parameter and two groups), one-way ANOVA (one parameter and more than two groups) when data showed normal distribution as evaluated by the Shapiro-Wilk test, or nonparametric Mann-Withney (two groups) or Kruskal-Wallis test (more than two groups). For analyses of at least two parameters, two-way ANOVA was used. For the fertility rate and ovaries containing multinucleated giant cells, a chi-square test was applicated. Data are shown as means ± SEM. A *P* value < 0.05 was considered as significant.

## Results

### Effects of reducing minipubertal gonadotropin levels by a GnRHR antagonist on reproductive parameters

Our previous studies on mice of the C57BL/6J strain revealed a considerable increase in FSH levels at the time of minipuberty (François *et al*., 2017). We obtained similar results in mice of the Swiss strain, with maximal FSH levels reached around 14 dpn (⋍20,000 pg/mL), being then 40-45-fold higher than those in peripubertal females at 28 dpn (Fig. 1A). To investigate the role of minipuberty on reproductive parameters, we used the same pharmacological strategy to decrease the levels of gonadotropins specifically at the time of minipuberty, as described before (François *et al*., 2017; Devillers *et al*., 2019). The treatment by the GnRHR antagonist (Ganirelix®) for seven days led to a 70-75 % decrease in circulating FSH levels at 14 dpn, close to what we previously obtained in C57BL/6J mice (François *et al*., 2017). Importantly, on the days after the last injection performed at 16 dpn, FSH levels returned to similar levels to those observed in CTR females, as observed at 17 and 21 dpn (Fig. 1B).

**Figure 1:**
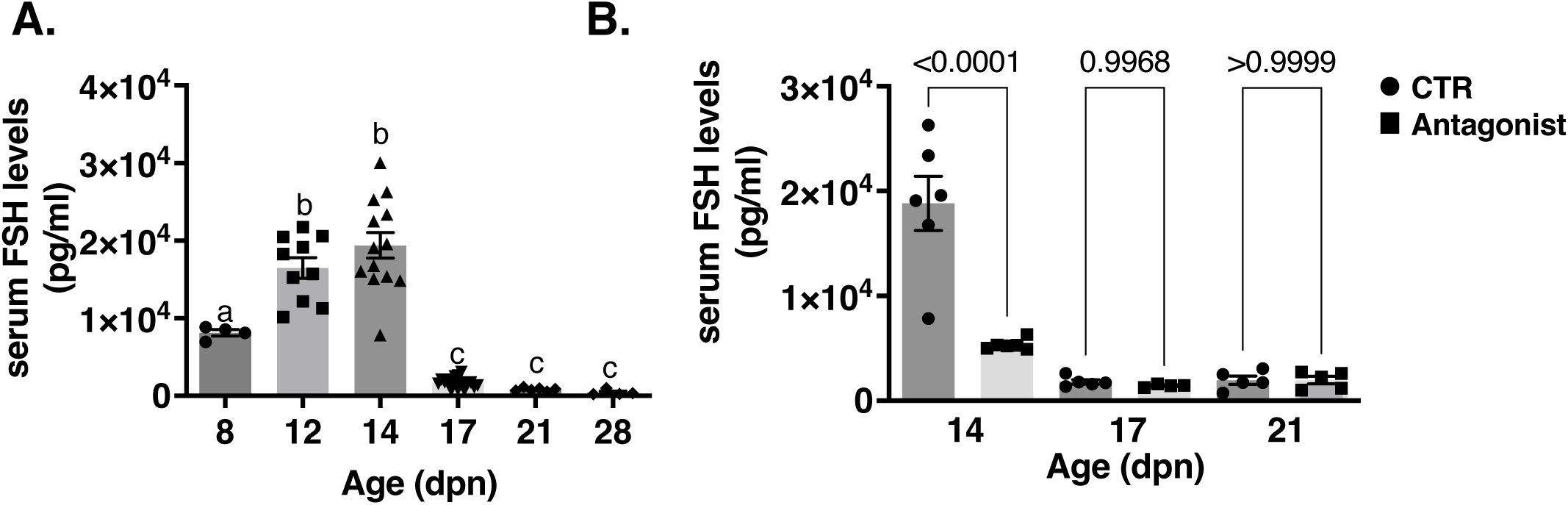
Determination of circulating FSH levels during the prepubertal period in Swiss mice and after treatment by the GnRHR antagonist during minipuberty. **(A)** Circulating levels of FSH measured by Luminex assay in wild-type mice at 8, 12, 14, 17, 21 and 28 dpn (4 to 12 samples per age). **(B)** Serum concentration of FSH measured by Luminex assay at 14, 17 and 21 dpn in CTR and GnRHR antagonist-treated mice (4 to 6 samples/age in CTR and antagonist-treated mice). In graphs, bars are the means ± SEM. Data were analyzed using a one-way ANOVA test **(A)** and a Mann-Whitney test **(B)**. Distinct letters indicate significant differences between groups, with *P*<0.05.

We evaluated the onset of puberty by analyzing on vaginal smears parameters used to define this event: vaginal opening, corresponding to the E2-dependent rupture of the filaments closing the vagina, as well as the occurrence of first estrus and first diestrus 2 (Fig. 2A). The observation of vaginal opening, spanning in our mouse colony over 21 and 32 dpn (mean age: 25.6 ± 0.4 dpn for CTR and 26.7 ± 0.4 dpn for antagonist-treated mice, n=28 mice/group) showed no significant difference between the two groups. We also found no difference in the occurrence of first estrus (mean age: 27.6 ± 0.4 dpn for CTR and 28.3 ± 0.5 for antagonist-treated mice, n=28 mice/group) and first diestrus 2 (mean age: 30.8 ± 0.3 dpn for CTR and 31.4 ± 0.5 for antagonist-treated mice, n=28 mice/group). Taken together, these results suggest that the significant reduction in FSH levels at minipuberty had no impact on the initiation of puberty in mice.

**Figure 2:**
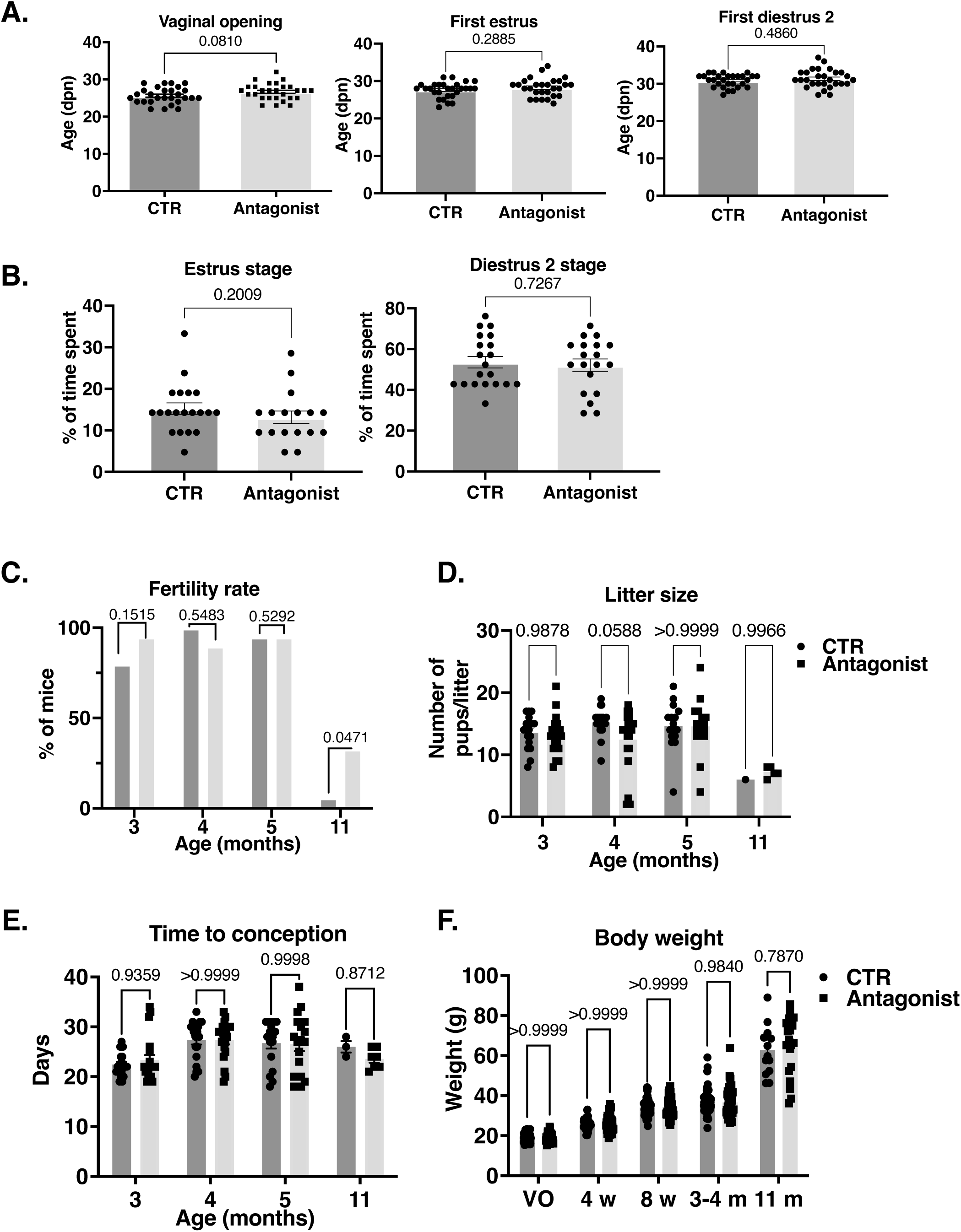
Reproductive parameters in mice with reduced minipubertal gonadotropin levels. **(A)** Determination of the age of vaginal opening, first estrus and first diestrus 2, by daily morning vaginal smears (28 females/group). **(B)** Percentage of time spent in estrus and diestrus 2 in 3- to 4-month-old females on 21 consecutive days of vaginal smears (20 females for CTR and 17 for antagonist-treated mice). **(C)** Fertility rate based on the percentage of fertile mice at 3, 4, 5 and 11 months (n=20 females/group at 3,4 and 5 months; 17 females for CTR and 15 females for antagonist-treated mice at 11 months). **(D)** Litter size at 3, 4, 5 and 11 months based on the number of pups seen at birth (20 females/group at 3,4 and 5 months; 17 females for CTR and 15 females for antagonist-treated mice at 11 months). **(E)** Time to conception, corresponding to the time between mating and parturition (20 CTR and 19 antagonist-treated females at 3,4 and 5 months; 17 females for CTR and 15 females for antagonist-treated mice at 11 months). **(F)** Body weights at the time of vaginal opening (VO), 4, 8 weeks and 3-4 and 11 months (18 to 62 females for CTR and 16 to 61 for antagonist-treated mice). In graphs, bars are the means ± SEM. Data were analyzed using Student t-test or Mann-Whitney test **(A, B)**, a Chi-square test **(C)** and a two-way ANOVA test **(D-F)**.

To assess whether the treatment altered estrous cyclicity, we performed vaginal smears for 21 consecutive days in adult females aged 3-4 months old (Fig. 2B). Our results showed that there was no difference in the percentage of time spent in estrus (15.2% versus 13.2% in CTR mice; *P=*0.2009) and diestrus 2 (53.5% versus 52.7% in CTR mice; *P=*0.8283) between the two groups, with considerable variability between females within the same group.

To further evaluate female reproductive parameters, we also analyzed the fertility of these mice (Fig. 2C). Females were mated with males as soon as the beginning of reproductive life, i.e., at 3 months of age, up to 11 months. We observed no difference in the percentage of fertile females between the two groups at either 3, 4 or 5 months of age, with a high breeding performance reaching 85 to 100% during this period (Fig. 2C). There was also no significant difference in the number of pups per litter between the two groups or in the delay of conception (i.e., in the time between mating with the male and pup delivery), which was comprised between 18 and 38 days in the two groups (Fig. 2D and E). When these mice were mated at 11 months of age, we observed that only 6% of CTR mice could still have pups, indicating that they have already undergone reproductive senescence at this age (Fig. 2C). This contrasted with the observation that 33% of antagonist-treated mice could still deliver pups (*P=*0.0471). Among the females that were fertile at this stage, we did not observe any significant difference in the delay of conception (Fig. 2E). The number of pups was much smaller than at earlier studied ages (about 6 to 8 pups/female), and it did not differ between the two groups (Fig. 2D).

Monitoring of body weights at different stages (i.e., vaginal opening (VO), 4, 8 weeks, 3-4 months, and 11 months), showed that the treatment by the antagonist had no significant impact on this parameter (Fig. 2F).

Overall, these findings suggest that suppressing minipuberty in mice had no effect on puberty initiation and early reproductive life, but that it increased reproductive longevity in a fraction of middle-aged females.

### Effect of minipubertal GnRHR treatment on neuroendocrine markers of reproduction

We next tested the hypothesis that increased reproductive lifespan in antagonist-treated mice could be associated with alterations in neuroendocrine aging related to hypothalamic control of reproduction. We, thus, measured the basal levels of gonadotropins in mice used for mating studies, including those that were infertile at middle-age. When we compared basal LH levels between females at 4 (young) and 11 (middle-age) months, we observed that they became significantly higher in middle-aged mice of the CTR group than in young mice of both groups (mean levels in the CTR group: 1027 ± 226.3 pg/mL at 11 months versus 294 ± 71.75 pg/mL at 4 months; *P*=0.0091) (Fig. 3A). In contrast, basal LH levels of middle-aged females treated with the GnRHR antagonist at minipuberty were similar to those observed in young females, and much lower than those of middle-aged CTR females (Fig. 3A). When we measured basal FSH levels, we observed no significant differences in its concentrations between young and middle-aged mice in both groups, although they tended to be lower in antagonist-treated mice than in CTR mice at 11 months (Fig. 3A).

**Figure 3:**
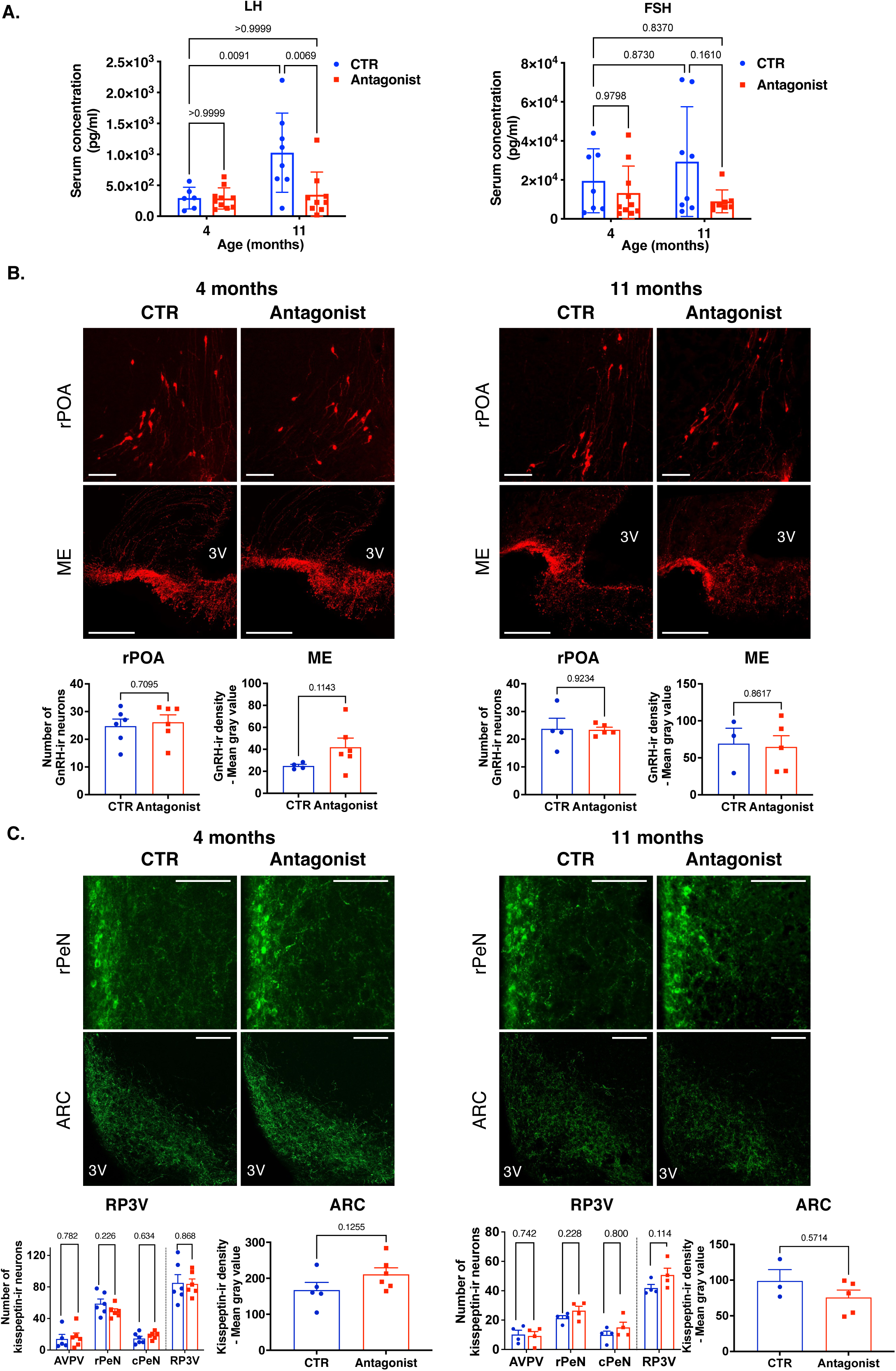
Analysis of the impact of reduced minipubertal gonadotropin levels on central determinants of reproductive function in young and middle-aged mice. **(A)** Serum concentrations of basal LH and FSH levels were measured by Luminex assay at 4- and 11-months in CTR and antagonist-treated mice (6 to 10 mice/group). **(B)** Representative images and quantification of the number of GnRH-immunoreactive (ir) neurons in the rostral preoptic area (rPOA) and mean density of GnRH immunoreactivity in the median eminence (ME) of 4- and 11-month-old females in diestrus (3 to 6 females/age and treatment group). **(C)** Representative images and quantification of the number of kisspeptin immune-reactive (ir) neurons in the rostral periventricular area of the third ventricle (RP3V) composed of the anteroventral periventricular (AVPV), rostral (rPeN) and caudal (cPeN) preoptic periventricular nuclei and mean density of kisspeptin immunoreactivity in the arcuate nucleus (ARC) of 4- and 11-month-old females in diestrus (n=3 to 6 females/ age and treatment group). 3V = third ventricle. Scale bar = 100µm. In graphs, bars are the means ± SEM. Data were analyzed using a one-way ANOVA test **(A)**, Student t-test (4 months) or Mann-Whitney tests (11 months) **(B, C)**.

We sought that reduced gonadotropin levels at minipuberty may have effects on GnRH and/or kisspeptin immunoreactivity (ir) during reproductive life to impact its longevity. We, thus, performed immunofluorescence studies on brain tissue sections in young (4 months-old) and middle-age (11 months-old) mice in diestrus. We found no significant differences between CTR and treated females in the number of GnRH neurons in the rostral preoptic area (rPOA) and mean GnRH-ir density in the median eminence (ME) at any age (Fig. 3B). We analyzed the number of kisspeptin cells in the anteroventral periventricular (AVPV), rostral (rPeN) and caudal (cPeN) preoptic periventricular nuclei, commonly referred to as the RP3V, as well as the mean density of kisspeptin-ir in the ARC (Fig. 3C). There was no apparent effect of the treatment on the number of kisspeptin cells in the AVPV, rPeN or cPeN, and overall, in the RP3V, at any studied age, although their number tended to increase in antagonist-treated mice compared with CTR at 11 months (mean: 41.9 versus 50.8, *P*=0.114). There was no significant effect on the mean kisspeptin-ir density in the ARC between the two groups, regardless of the age studied (Fig. 3C).

### Effect of minipubertal GnRHR antagonist treatment on follicular growth and atresia in the ovaries of middle-aged females

To determine whether antagonist-treated mice exhibited higher reproductive capabilities at 11 months due to improved follicular endowment, we performed morphometric analyzes on the ovaries by automatic counting of follicles and corpora lutea (CL). Our *in situ* observations revealed that the ovaries of both groups contained follicles at different stages, and CL (Fig. 4A). However, the number of primordial follicles was extremely reduced at this stage. Interestingly, we observed that their number was significantly reduced by about 80% in antagonist-treated mice as compared with CTR (mean: 11.7 ± 3.5 versus 2.2 ± 1.0 in antagonist-treated mice; *P=*0.0355) (Fig. 4B). The number of follicles at primary and secondary stages was also lower in antagonist-treated mice than in CTR mice, with an observed decrease of 63% and 50%, respectively (*P*= 0.0296 and 0.0381, respectively) (Fig. 4B). In contrast, there was no difference in the number of tertiary follicles between the two groups (Fig. 4B). There was no significant difference in the number of atretic follicles of any category between the two groups, although antagonist-treated mice had about 40% fewer atretic follicles at the secondary stage than CTR mice (mean: 18.0 ± 3.0 in CTR versus 9.83 ± 2.44 in antagonist-treated mice; *P=*0.0631) (Fig. 4C). Since the presence of corpora lutea (CL) indicates ovulation, we also investigated their presence and number. As there were CL of different sizes, we classified them in category 1 (small: regressed), 2 (intermediary: formed one or more cycles before) and 3 (large: formed during the current cycle). We observed no difference in the number of any category between the two groups (Fig. 4D). In addition, we rarely observed oocytes trapped within CL, a feature linked to defective ovulation, in either CTR or antagonist-treated ovaries (Mara *et al*., 2020). Determination of circulating progesterone levels, which is mainly produced by CL, did not reveal any difference between the two groups, suggesting that luteal function was comparable (Fig. 4E). Taken together, these results suggest that despite their enhanced reproductive capacity, antagonist-treated mice exhibited greater follicular depletion than CTR mice but maintained ovulation.

**Figure 4:**
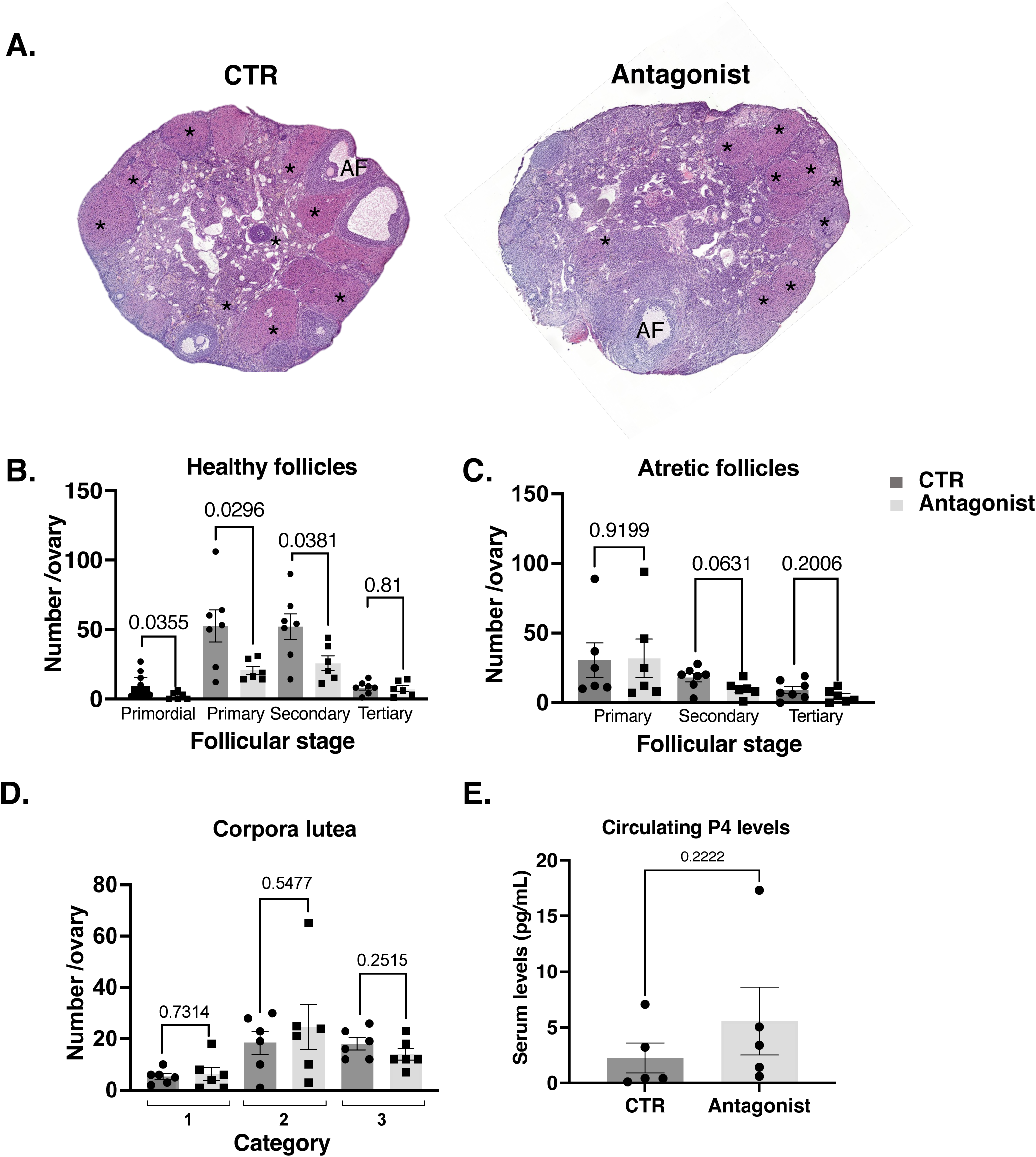
Impact of the GnRHR treatment on follicular and corpora lutea content in middle-aged mice. **(A)** Histological ovarian sections of middle-age ovaries from CTR and antagonist-treated mice, showing corpora lutea (*) and antral follicles (AF). **(B-C)** Morphometric analysis of the number of healthy **(B)** and atretic follicles **(C)** in each category at 11 months (6 to 7 ovaries from different females/group). **(D)** Morphometric analysis of the number of corpora lutea (CL), which were classified in 3 categories according to their size and the presence or not of apoptotic luteal cells (regressing: type 1; formed one or more cycles before: type 2; formed during the current cycle: type 3) (6 to 7 ovaries from different females/group). **(E)** Serum concentrations of progesterone levels were measured by GC-MS (5 samples from different females/group). In graphs, bars are the means ± SEM. Data were analyzed using Student t-test or Mann-Whitney test **(B-E)**.

### Effect of the minipubertal GnRHR antagonist treatment on key endocrine markers of ovarian function in middle-aged females

We then investigated in middle-aged mice the effect of the antagonist treatment on the levels of key follicular markers associated with endocrine function: anti-Mullerian hormone, (AMH), the aromatase enzyme converting androgens into estrogens *(Cyp19a1)*, Inhibin ꞵA and ꞵB subunits *(Inhba* and *Inhbb)*, and the receptors for FSH (*Fshr*) and LH (*Lhcgr*) (Fig. 5A). AMH expression is found primarily in granulosa cells of healthy growing follicles from the primary to the antral follicle stage and is low or absent in mural granulosa cells and atretic follicles (Baarends *et al*., 1995). *Cyp19a1*, *Inhba, Inhbb and Fshr* are mainly expressed by granulosa cells of tertiary follicles, and *Lhcgr* is expressed in thecal cells of growing follicles and in granulosa cells of advanced tertiary follicles (Richards and Pangas, 2010). Circulating AMH levels and the relative intra-ovarian abundance of *Amh* transcripts were significantly lower in antagonist-treated mice than in CTR mice, consistent with the observed reduction in primary and secondary follicle numbers (Fig. 4B, 5B and C). In contrast, the relative intra-ovarian expression of *Cyp19a1*, *Inhba*, *Inhbb and Fshr*, and that of *Lhcgr*, was not significantly affected by the antagonist treatment (Fig. 5D-H). Overall, our results suggest that the increase in follicular depletion observed in antagonist-treated mice was essentially associated with a decrease in AMH synthesis by the ovary, without apparent alteration in the other endocrine parameters studied.

**Figure 5:**
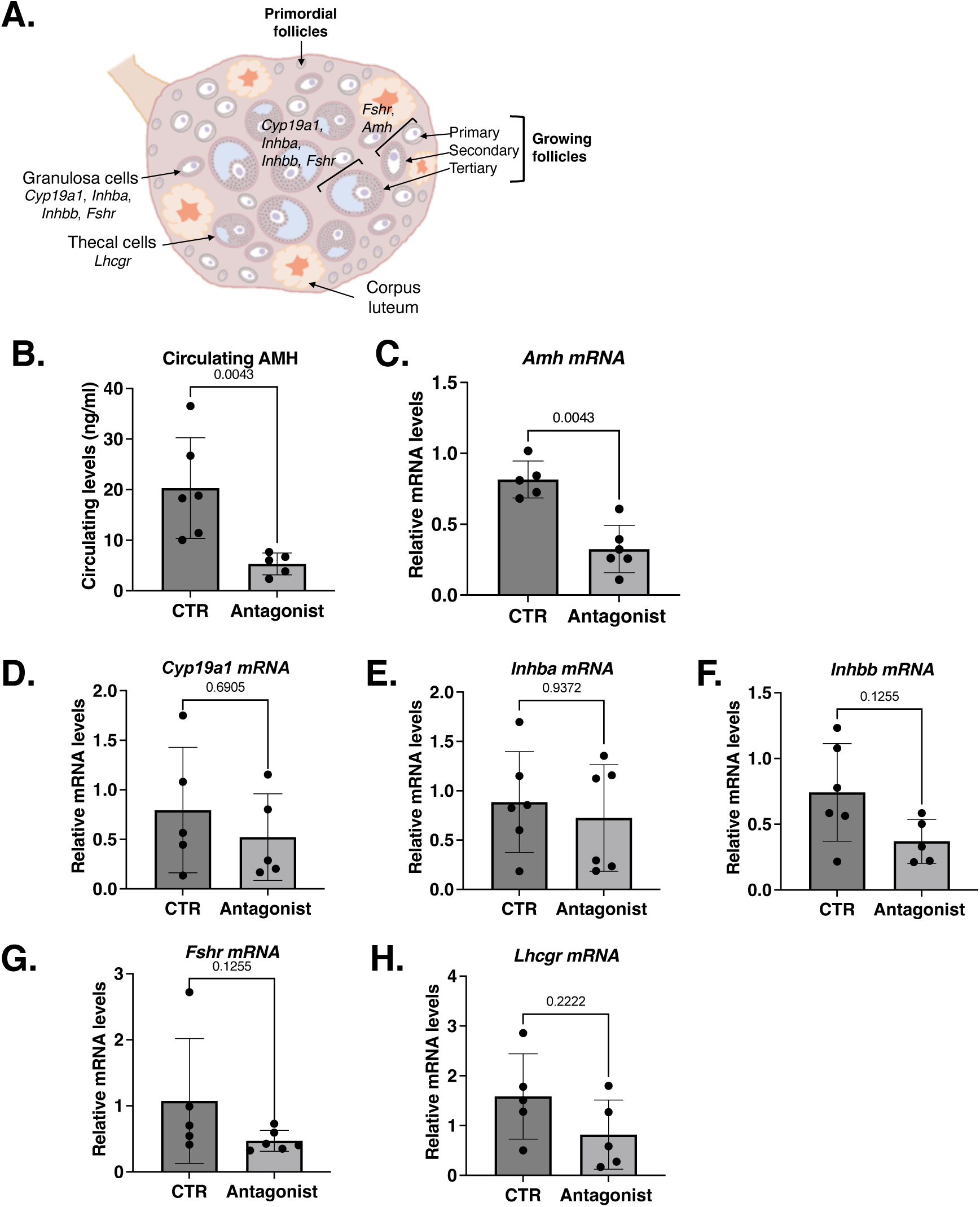
Analysis of ovarian function in middle-age mice treated with the GnRHR antagonist at minipuberty. **(A)** Schematic representation of key markers of endocrine activity in the ovary. Shown are the androgen-converting enzyme *Cyp19a1* aromatase, the inhibin beta subunits (*Inhba*, *Inhbb*), *Amh*, and *Fshr*, which are all expressed in granulosa cells of growing follicles and *Lhcgr* (encoding the LH receptor) expressed in thecal cells. **(B)** Serum concentration of AMH was measured by ELISA assay at 11 months (5 to 6 samples from CTR and antagonist-treated mice). **(C-H)** Relative intra-ovarian abundance of *Amh*, *Cyp19a1*, *Inhba, Inhbb*, *Fshr* and *Lhcgr* transcripts in CTR and antagonist-treated mice aged 11 months determined by quantitative real-time RT-PCR and normalized to the mRNA levels of *Hprt* (5 to 6 ovaries from different females/group). In graphs, bars are the means ± SEM. Data were analyzed using a non-parametric Student t-test **(B-H)**.

### Effect of reduced minipubertal gonadotropin levels on markers of ovarian aging in middle-aged mice

To better understand the effect of GnRHR treatment administered during minipuberty on the ovaries of middle-aged mice, we analyzed characteristics of ovarian aging: oocytes-depleted follicles originating from small follicles that have lost their oocytes (oocyte-depleted follicles, abbreviated as ODF) (Crumeyrolle-Arias and Aschheim, 1981; Guigon *et al*., 2005) and multinucleated giant cells resulting from the fusion of several macrophages (Zhang *et al*., 2020; Camaioni *et al*., 2022). Both ODF and multinucleated giant cells were visible in the ovaries of CTR and antagonist-treated mice at 11 months (Fig. 6A-C). There appeared to be less ODF in antagonist-treated mice compared with CTR mice, but this reduction was not significant (Fig. 6B). Additionally, there was a greater percentage of multinucleated giant cells in CTR ovaries than in antagonist-treated ovaries, but again, this was not significant (71% in CTR versus 33% in antagonist-treated mice; n=6 to 7 ovaries/group; *P=*0.1696) (Fig. 6C).

**Figure 6:**
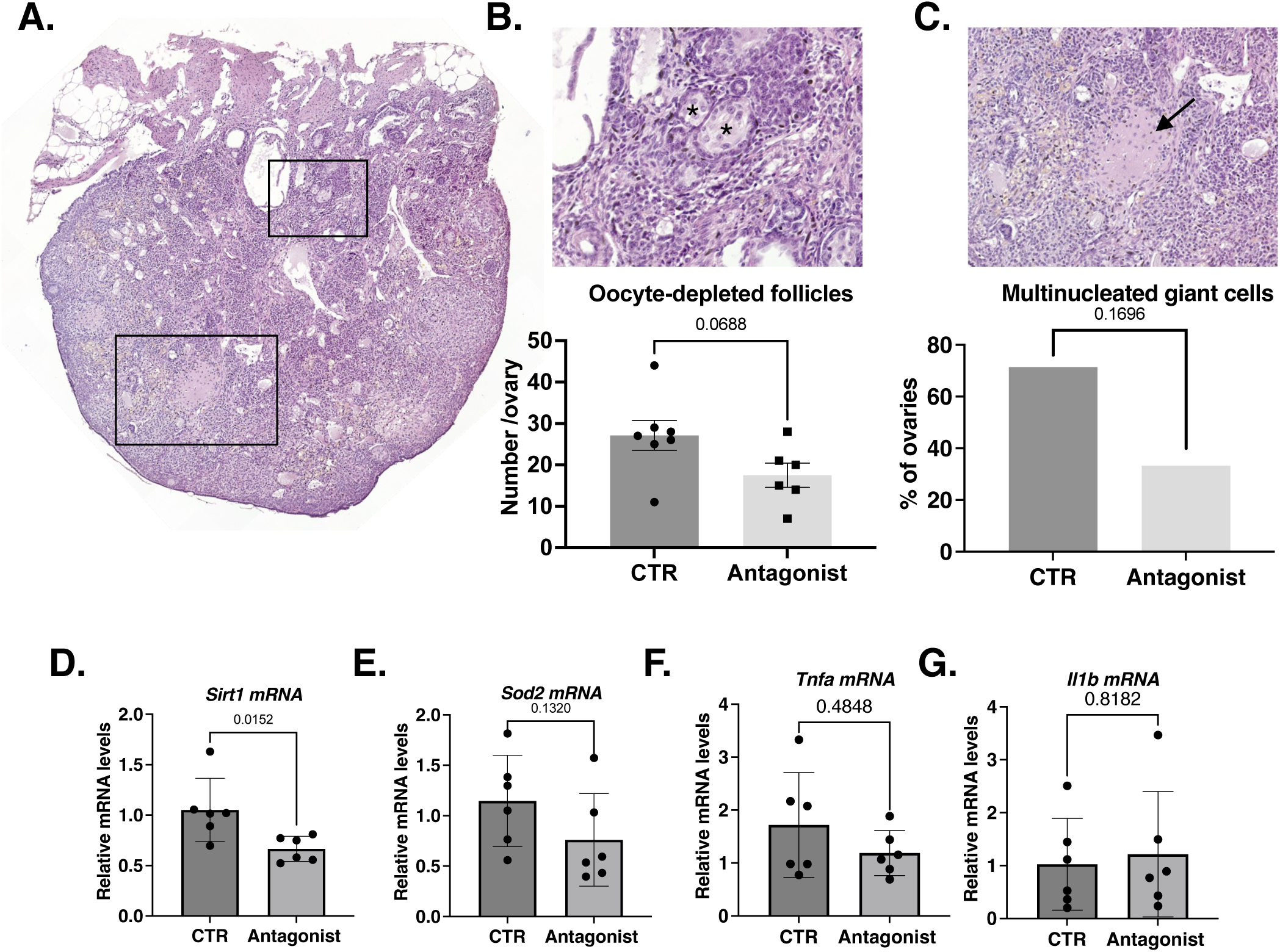
Ovarian aging and inflammation in middle-aged mice treated with the GnRHR antagonist at minipuberty. **(A)** Histological ovarian sections of middle-age ovaries from CTR and antagonist-treated mice showing typical signs of aging, i.e., oocyte-depleted follicles **(B)** and multinucleated giant cells **(C)**. Morphometric analyses of oocytes-depleted follicles at 11 months (6 to 7 ovaries from different females/group). **(C)** Percentage of ovaries with multinucleated giant cells at 11 months (6 to 7 ovaries from different females/group). **(D-G)** Relative intra-ovarian abundance of *Sirt1*, *Sod2*, *Tnfa* and *Il1b* transcripts in the ovaries of 11 months for CTR and antagonist-treated mice determined by quantitative real-time RT-PCR and normalized to the mRNA levels of *Hprt* (6 ovaries from different females/group). In graphs, bars are the means ± SEM. Data were analyzed using Student t-tests (B, D-G) and Chi-square test (C).

We also investigated the contribution of oxidative stress, a process which is involved in ovarian aging, by analyzing the gene expression of sirtuin 1 (*Sirt1)* and manganese superoxide dismutase *Sod2* (Liu *et al*., 2013). Indeed, *Sirt1* expression increases in aging ovary, possibly to adapt its response to increasing oxidative stress, in particular *via* induction of *Sod2* expression (Di Emidio *et al*., 2014; Tatone *et al*., 2018). When we analyzed the expression of these two genes, we observed that the relative intra-ovarian abundance of *Sirt1* transcripts was significantly lower in antagonist-treated mice than in CTR mice and that of *Sod2* tended to decrease, but this was not significant (Fig. 6D and E). Age-related follicular depletion has also been reported to be associated with increased inflammation and elevated expression of certain markers such as tumor necrosis factor alpha (*Tnfa*) and interleukin 1 beta (*Il1b*) (Navarro-Pando *et al*., 2021). However, our results did not show any difference in their relative expression levels between the two groups (Fig. 6F and G). All these data suggest that higher follicular depletion observed in antagonist-treated mice occurred without obvious acceleration of ovarian aging but surprisingly with a seemingly reduced response to oxidative stress.

## Discussion

Although it is clearly established that developmental events occurring during fetal life are essential for subsequent female reproductive function, the possible role of the postnatal period of minipuberty remains to be established. In the present study, we provide evidence, using mice as an experimental model, that minipuberty contributes to a major aspect of fertility, notably by regulating its duration.

Indeed, we observed that the reduction in gonadotropin levels by GnRHR antagonist treatment during minipuberty resulted in a longer reproductive lifespan at middle age, since approximately 33% of them were able to have pups at 11 months and only 6% of CTR females. This increase in reproductive lifespan was not accompanied by a change in the age of initiation of puberty or in the reproductive capacity and estrous cyclicity in earlier life, since these parameters were comparable between the two groups of mice.

By investigating the mechanism underlying reproductive lifespan in our model, we discovered that middle-aged mice treated by the GnRHR antagonist at minipuberty maintained basal LH levels equivalent to those observed in young adult mice, unlike their age-matched CTR displaying higher basal LH levels associated with reproductive senescence (Parkening *et al*., 1980). The fact that this occurs before any change in ovarian activity has led to propose that the advent of reproductive senescence results mainly from neural aging in rodents (Belisle *et al*., 1990; Kermath and Gore, 2012; Guo and Pankhurst, 2020). The increase in basal LH levels observed in middle-aged rodents would reflect the alteration in GnRH pulsatility due to decreased estradiol receptivity in the hypothalamus (Yaghmaie *et al*., 2010). Indeed, in our study, we observed that the measurable differences in circulating LH levels between CTR and GnRHR antagonist-treated mice was not accompanied by the modification in the expression of the genes encoding aromatase (the enzyme responsible for the production of E2) and Inhibin beta A and B subunits contributing to Inhibin/Activin A and B, suggesting that the levels of these hormones regulating the activity of the hypothalamus and the pituitary remained unaltered. Although the mechanisms underlying hypothalamic aging are unknown, some studies describe a decrease in the number of kisspeptin neurons (Kermath *et al*., 2014). Our immunofluorescence experiments showed no difference in the number of GnRH neurons in rPOA and GnRH-ir density in the ME between the two groups, regardless of age. In addition, there was no change in the mean density of kisspeptin-ir in the ARC or in the number of kisspeptin neurons in the RP3V, although this latter tended to increase in the GnRHR antagonist-treated group when compared to middle-age CTR. It is, however, important to stress that quantification of kisspeptin-immunoreactivity was performed in female mice at the diestrus stage to avoid the great variability in estrogen levels during the follicular phase. Further studies should address in details the expression of kisspeptin at proestrus and estrus stages in female mice minipubertally treated with GnRHR antagonist. Alternatively, the pulsatility of GnRH neurons is driven by a complex cellular network, including astrocytes which are recruited after birth by GnRH neurons (Pellegrino *et al*., 2021). Astrocytes could contribute to aging-related reproductive function in female mice (Dai *et al*., 2020). Further studies are needed to understand the mechanisms involved in the maintenance of LH levels of young adult females in antagonist-treated mice at middle-age, and to establish whether it resulted from delayed neural aging by modulation of hypothalamic networks regulating reproduction.

Paradoxically, the prolonged reproductive lifespan in middle-aged mice treated with the GnRHR antagonist during minipuberty occurred despite accelerated follicular depletion, as shown by lower numbers of primordial, primary, and secondary follicles per compared to control mice. Using AMH as a marker of follicle growth consistently revealed its lower circulating levels and relative intra-ovarian transcript abundance in antagonist-treated females compared to controls. Furthermore, the relative abundance of *Cyp19a1* and *Inhba* transcripts, which are primarily expressed by granulosa cells of large antral follicles, was not different between the two groups, consistent with their comparable number of follicles in this category. The similar numbers of corpora lutea and circulating progesterone levels in the two groups suggests that terminal maturation of follicles up to ovulation was unaltered in antagonist-treated mice at middle-age, despite lower numbers of primordial and early growing follicles than controls. Therefore, reproductive performance was not correlated to the size of the follicular reserve, as already observed in other models, such as *Amh^-/-^* mice which had a similar age of onset of infertility despite lower levels of primordial follicles than *Amh^+/+^* mice (Guo and Pankhurst, 2020). Our data together with others further highlight the idea that the primary mechanism of reproductive senescence in rodents results from neuroendocrine aging rather than on follicular depletion, as previously discussed (Kermath and Gore, 2012).

The mechanisms underlying the decreased number of primordial and early growing follicles in the ovaries of middle-aged females treated with the GnRHR antagonist during minipuberty are currently unclear. The possibility of accelerated ovarian aging is unlikely since it was not associated with an increase in structures typically seen in aging ovaries, such as ODF and multi-nucleated giant cells, or in increased expression of markers of oxidative stress and inflammation such as *Sirt1*, *Sod2*, *Tnfa* and *Ilb1* (Di Emidio *et al*., 2014; Tatone *et al*., 2018). Rather, we suspect that it resulted from the modification of follicular dynamics during minipuberty induced by the reduction in gonadotropin levels and the possible alteration in the levels of AMH, which are key regulators of folliculogenesis (Richards and Pangas, 2010; di Clemente *et al*., 2021; Zhou *et al*., 2022).

In conclusion, the current study reports an unsuspected role of gonadotropins during minipuberty acting on the gonadotrope axis resulting in long-term effects on fertility. Although the mechanisms by which gonadotropins exert this action remain to be explored, it is possible that they involve ovarian hormones, such as estradiol, AMH and/or androgens, which are regulated by gonadotropins and display receptors in the hypothalamus and the ovary during this period (Brock *et al*., 2015; Cimino *et al*., 2016; François *et al*., 2017; Devillers *et al*., 2019, 2023; Watanabe *et al*., 2023). These hormones may regulate the maturation of hypothalamic networks related to reproduction as well as follicular dynamics in the ovary. Our study raises concerns about the possible consequences on reproductive health of disturbances in the activity of minipubertal gonadotrope axis, as observed during premature birth (Kuiri-Hänninen *et al*., 2011), or which may occur following exposure to endocrine disruptors.

## Acknowledgments

The authors wish to thank the staff members of the core animal facility Buffon of University Paris Cité (Paris, France). They acknowledge the technical assistance of M. Surenaud for gonadotropin assays (Hôpital Henri Mondor, Créteil, France). They also thank F. Caud (Université Paris-Saclay), F. Clément (INRIA) and R. Yvinec (INRAE).

## Authors’ roles

CJG, MMD, LN & SMK designed the experiments. MC, MMD, EA, RC, FS, DQ, CHP, FG, LN, and CJG performed and analyzed the experiments. MC, LN, RC and CJG wrote the initial draft. CJG, MC and LN prepared the figures. CG, FG, LN and SMK edited the article. All co-authors approved the manuscript.

