## Supplementary Table 1 for "Minipuberty regulates reproductive lifespan and ovarian follicular loss in a mouse model with reduced minipubertal gonadotropin levels"

**Supplementary Table 1: GC/MS analytical control validation**.

| **Accuracy** (%) | **Analytes** | **Precursor ion analyte (GC-MS/MS) / IS*** (m/z) | **Range** (pg) | **Mean** (pg/ml) ***Intra- & Inter-assay CVs*** *(%)* | | | |
|---|---|---|---|---|---|---|---|
| | | | | **LLOQ** Mean Intra- & Inter assay CVs | **Low QC** Mean Intra- & Inter assay CVs | **Middle QC** Mean Intra- & Inter assay CVs | **High QC** Mean Intra- & Inter assay CVs |
| 94 - 110 | Prog / Prog-d9 | 510.25>510.20<br>510.25>147.20<br>510.25>495.20<br>519.25>519.20*<br>519.25>147.20* | 20 - 4 860 | 19.6 | 248.8 | 499.1 | 998.3 |
| | | | | *17.4 - 21.3* | *5.5 - 7.4* | *3.3 - 7.8* | *3.5 - 6.8* |

*LLOQ : low limit of quantification; QC : quality control*
